# Single-nucleus RNA-seq reveals disrupted cell maturation by chorioamnionitis in the preterm cerebellum of nonhuman primates

**DOI:** 10.1101/2022.07.12.499720

**Authors:** Josef Newman, Xiaoying Tong, April Tan, Vanessa Nunes De Paiva, Pietro Presicce, Paranthaman S Kannan, Kevin Williams, Andreas Damianos, Marione Tamase Newsam, Merline Benny, Shu Wu, Karen Young, Lisa A. Miller, Suhas G Kallapur, Claire A Chougnet, Alan H Jobe, Roberta Brambilla, Augusto F. Schmidt

## Abstract

Preterm birth is often associated with chorioamnionitis and increased risk of neurodevelopmental disorders. Preterm birth also leads to cerebellar underdevelopment but the mechanisms of disrupted cerebellar development in preterm infants are little understood. Here, we leveraged single-nuclei RNA-sequencing of the cerebellum in a rhesus macaque model of chorioamnionitis and preterm birth, to show that chorioamnionitis leads to Purkinje cell loss and disrupted maturation of granule cells and oligodendrocytes in the fetal cerebellum at late gestation. Purkinje cell loss is accompanied by decreased sonic hedgehog signaling from Purkinje cells to granule cells, which show an accelerated maturation. Chorioamnionitis also accelerated pre-oligodendrocyte maturation into myelinating oligodendrocytes, confirmed by increased expression of myelin basic protein in the cerebellum of chorioamnionitis-exposed fetuses. These findings are consistent with reported histopathological findings in individuals with autism and suggest a novel mechanism through which perinatal inflammation contributes to neurodevelopmental disorders.

## Introduction

The incidence of neurodevelopmental disorders is increasing among children, in part due to improved survival of at-risk newborns, such as extreme preterm infants^1,2^. Preterm infants who are exposed to inflammation prenatally, most commonly due to inflammation of the amniotic membranes^3^, or chorioamnionitis, are at particularly high risk of neurodevelopmental disorders, including autism spectrum disorders (ASD)^4,5^. While much of the research on brain injury and neurodevelopmental outcomes in preterm infants has focused on motor disabilities associated with supratentorial injury, disruption of cerebellar development is often seen in preterm infants, particularly underdevelopment^6,7^. Our understanding of the cerebellar function has shifted from a largely motor and coordination role to a broader understanding of its importance in the regulation of cognitive function, including attention, memory, and executive functioning^8,9^. In fact, the cerebellum is one of the most consistently abnormal brain areas in individuals with autism^10^. The cerebellum undergoes rapid expansion in the third trimester of gestation with peak proliferation of the granule cells, the most abundant neuron in the CNS, in the external granule layer between 26 and 32 weeks before migration to the internal granule layer^11^. Insults associated with preterm birth take place during the critical time of neuronal expansion of the cerebellum. Reduction of cerebellar volume among preterm infants has been shown to persist at least into adolescence and occurs even in the absence of prior injury being detected on brain imaging^12–14^. These findings suggest that events associated with prematurity and inflammation lead to impaired development of the cerebellum. Considering the importance of the cerebellum and of prenatal inflammation to the pathogenesis of neurodevelopmental disorders^15^, such as ASD, and their association with preterm birth^16,17^, our knowledge of the effects of chorioamnionitis to the cerebellum in this critical stage of development and their potential contribution to the pathogenesis of neurodevelopmental disorders remains limited.

To fill this gap in knowledge, we performed single-nucleus RNA-sequencing of the cerebellum in a non-human primate model of preterm chorioamnionitis. We demonstrate that exposure to chorioamnionitis decreases the number of Purkinje cells in the cerebellum and impairs proliferation signaling from Purkinje cells to granule cells in the external granule layer by decreased sonic hedgehog (SHH) signaling. Chorioamnionitis also led to early maturation and myelination of oligodendrocytes in the cerebellum. These findings are consistent with histopathological findings of individuals with autism and of autopsy reports of preterm infants^10,18,19^. Our findings provide new insight into the mechanisms through which inflammation contributes to the disrupts brain development in preterm infants.

## Results

### Intra-amniotic LPS induces persistent fetal inflammation after 5 days

To model chorioamnionitis in a preterm nonhuman primate we performed ultrasound-guided intra-amniotic (IA) injection of 1mg of LPS (E. coli O55:B5) on gestational day 127 (80% gestation, term is 165 days) (**Figure 1a, supplemental table 1**). This model has been previously shown to induce inflammation at the maternal-fetal interface, fetal systemic inflammatory response, and fetal neuroinflammation up to 48 hours after injection ^20,21^. Flow cytometry of the chorio-decidua showed that, at 5 days after IA LPS injection, there was a predominantly neutrophilic infiltrate in the chorio-decidua (**Figure 1b, supplemental Figure 1**). Inflammatory infiltrate in the membranes was associated with elevation of IL-6, IL-17, and IL-1ra in the fetal plasma by multiplex ELISA, without elevations of cytokines in the maternal plasma at 5 days (**Figure 1c, supplemental figure 2**).

**Figure 1.**
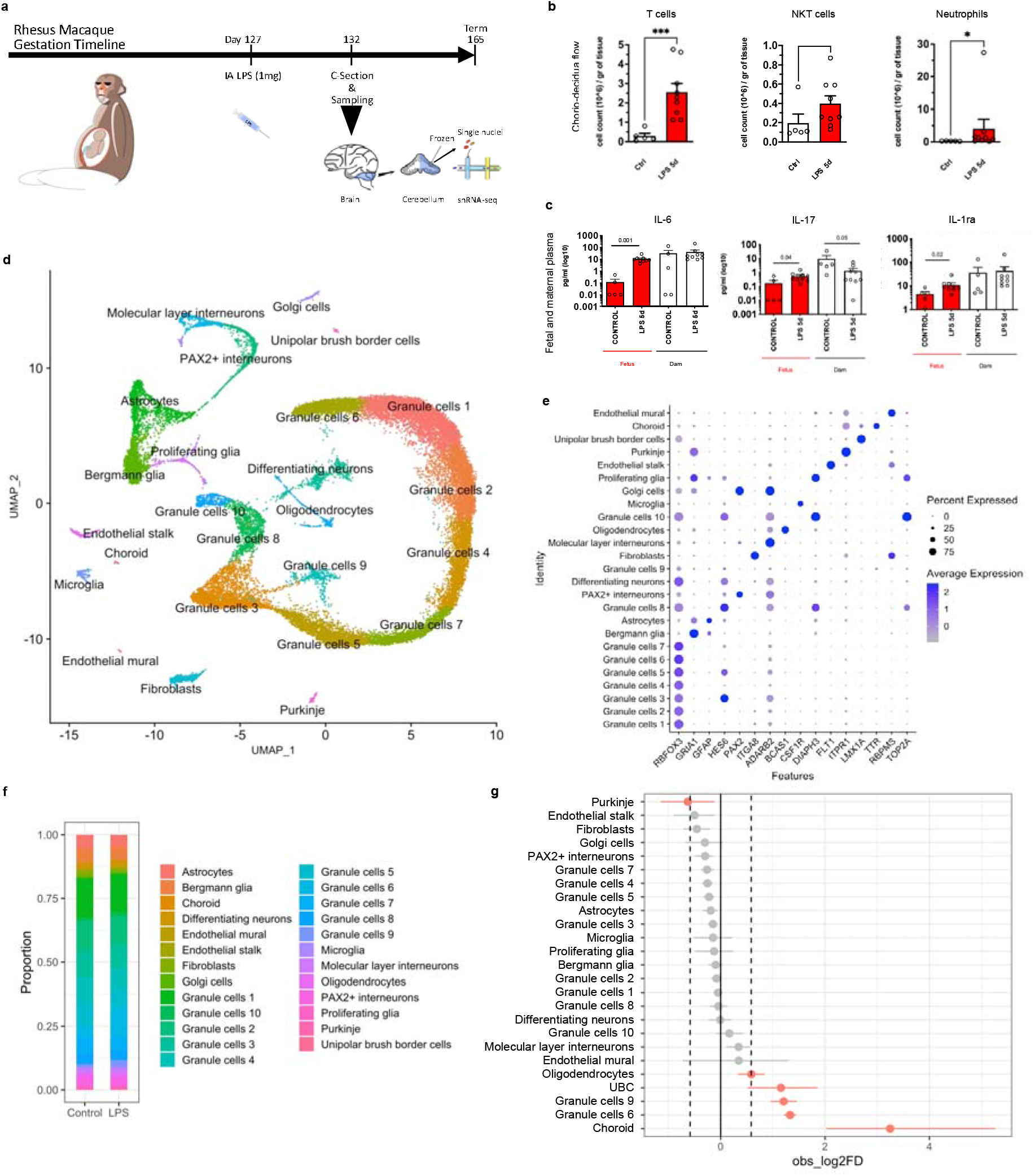
Intra-amniotic (IA) LPS induces persistent choriodecidual inflammation and fetal systemic inflammatory response similar to chorioamnionitis and leads to altered cerebellar cell composition. **a,** Experimental design. Rhesus macaques fetuses at 127 days of gestation (80% gestation) received ultrasound-guided intra-amniotic injections (IA) of 1mg of LPS from E. coli O55:B5 as a model of chorioamnionitis; n=7/group. **b,** Flow cytometry of the chorio-decidua cells showed increased neutrophils, T cells and NKT cells at 5 days in LPS animals (n=5/group). **c,** Multiplex ELISA for cytokines in the maternal and fetal serum from controls and LPS animals shows increased IL-6 as well as increased IL-17 and IL-1ra in the fetal plasma, with maternal increase in IL-17 only. (n=5/group). **d,** UMAP visualization of single-nuclei RNA-seq from 30,711 cerebellar nuclei. Clustering using the Seurat package revealed 24 unique cell clusters in the developing cerebellum; n= 2/group. **e,** Dotplot of top cell type markers based on the top differentially expressed genes in each cluster and on cell-specific and cell-enriched markers reported in the literature. **f,** Proportion of each identified cell cluster by condition showing predominance of granule cells in control and LPS animals. **g,** Differential proportion of cell populations in each cluster between control and LPS-exposed fetuses by permutation test. LPS decreased number of Purkinje cells and increased number of oligodendrocytes, unipolar brush border cells (UBC), and granule cell clusters 9 and 6.

### snRNA-seq reveals major developing cell types in the cerebellum at late gestation

To determine the effects of chorioamnionitis on the developing cerebellum we used unbiased high-throughput single-nucleus RNA-sequencing (snRNAseq) to examine transcriptional populations of the cerebellum from preterm rhesus macaque fetuses exposed to IA LPS or placebo. We then analyzed the transcriptome from 30,711 single nuclei (17,614 nuclei from 2 controls, 13097 nuclei from 2 chorioamnionitis) to an average depth of 25,192 to 45,124 reads per nuclei (**supplemental table 2**). Nuclei were clustered based on their expression profile and we identified 24 cell clusters that were annotated based on published cell markers^22–24^ (**Figure 1d, e supplemental table 3**). We then analyzed the cell type composition in the cerebellum of controls and LPS-exposed fetuses (**Figure 1f**). Permutation test^25^ demonstrated a decrease in the proportion of Purkinje cells and an increase in oligodendrocytes, unipolar brush border cells (UBC), choroid cells, and granule cell clusters 9 and 6 in LPS-exposed fetuses (**Figure 1g**), showing that chorioamnionitis alters the cell composition of the fetal cerebellum.

### Chorioamnionitis accelerates cerebellar granule cell maturation

Cerebellar granule cells are the most abundant neurons in the central nervous system and were the most abundant cell type identified in our dataset. Given the increased proportion of two granule cell clusters in LPS-exposed fetuses we re-clustered the granule cell populations for further analyses. Re-clustering of the granule cells identified seven distinct clusters based on their expression profile (**Figure 2a, b**). Analysis of the top cluster gene markers and established markers for neurons at different developmental stages identified granule cells at five stages of development in our dataset: proliferating granule cells, committing granule cells, migrating granule cells, maturing granule cells, and mature granule cells (**Figure 2c, d, supplemental table 4**). Differential expression analysis of gene expression between LPS-exposed and control cerebellum identified 317 differentially expressed genes using a threshold of log fold-change > 0.25 and FDR < 0.1 (**Figure 2e, supplemental table 5**). Overrepresentation analysis of differentially expressed genes was performed on ToppCluster^26^ for genes induced and suppressed in chorioamnionitis. Genes induced in granule cells of LPS-exposed fetuses were associated with synapse transmission and hippo signaling pathways, while suppressed gene were associated with cerebellum development, and Purkinje cell – granule cell precursor signaling involved in granule cell precursor proliferation (**Figure 2f**). To further explore the effect of chorioamnionitis on granule cell development we performed pseudotime analyses on Monocle3^27^ and Slingshot^28^ followed by differential topology between conditions^29^ (**Figure 2g**). Both pseudotime analysis methods showed an increased proportion of the most mature granule cells suggesting that chorioamnionitis accelerates cerebellar granule cell maturation.

**Figure 2.**
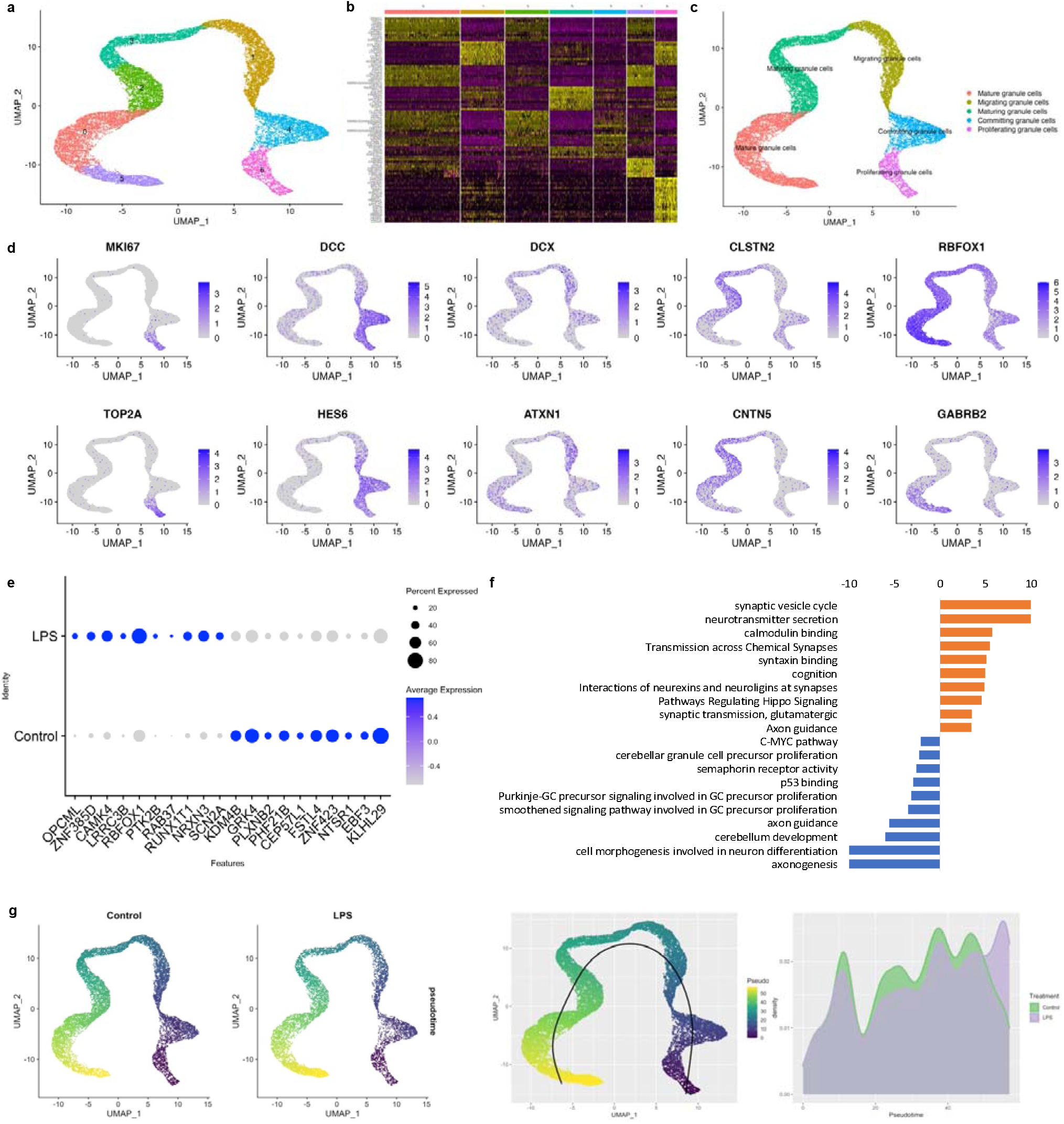
Prenatal inflammation disrupts cerebellar granule cell development increasing the number of mature granule cells. **a,** UMAP plot of re-clustered granule cells identified 7 distinct populations of granule cells in the developing cerebellum. **b,** Heatmap of the top 10 differentially expressed genes in each granule cell cluster identified shows similarities between some of the clusters which were regrouped for classification based on gene expression patterns. **c,** Regrouped clusters based on gene expression patterns shows the presence of granule cells at different developmental stages from proliferating to mature granule cells. **d,** Feature plots of identified cell markers for proliferative granule cells (MKI67 and TOP2A), granule cells exiting cell cycle and committing to differentiation (DCC and HES6), migrating granule cells (DCX and ATXN1), granule cells with increased mature cell markers and markers that the granule cells have exited the external granule layer (CNTN2 and CLSTN1), and mature granule cells (RBFOX3 and GABRB2). **e,** Dotplot of top genes differentially expressed in LPS-exposed fetuses vs. control in all clusters of the cerebellar granule cells. **f,** Overrepresentation analysis of differentially expressed genes in LPS-exposed fetuses vs controls. Genes induced by LPS were associated with biological processes for synapse function and hippo signaling pathway. Genes suppressed by LPS were associated with Purkinje-granule cell precursor signaling involved in granule cell precursor proliferation, cerebellum development, and cell morphogenesis involved in neuron differentiation. **g**, Pseudotime analysis on monocle3 and slingshot using the condiments package to identify differences in proportion of cells along pseudotime shows increased granule cell maturation LPS-exposed fetuses.

### Chorioamnionitis decreases the relative abundance of Purkinje cells in the developing cerebellum

Since IA LPS decreased the proportion of Purkinje cells, we sought to further determine the effect of chorioamnionitis on Purkinje cell numbers and gene expression. Sub-setting and re-clustering of the Purkinje cells revealed 2 clusters (**Figure 3a**) with distinct cell markers (**Figure 3b, supplemental table 4**). Overrepresentation analysis for Kegg pathways of genes differentially expressed in cluster 1 compared to cluster 0 of Purkinje cells showed that genes in cluster 0 were associated with Hippo signaling, cellular senescence, and SHH signaling while genes in cluster 1 were associated with glutamatergic synapse, Rap1 signaling pathway, and Ras signaling pathway, suggesting distinct developmental functions of these subpopulations (**Figure 3c**). Differential gene expression of genes expressed in Purkinje cells in LPS-exposed compared to control fetuses resulted in 302 regulated genes (**figure 3d, supplemental table 6**). Pathway overrepresentation analysis of genes differentially expressed in LPS-exposed fetuses compared to control showed that genes suppressed by chorioamnionitis were associated with cholinergic synapse, dopaminergic synapse, and Wnt signaling, and genes induced by LPS were associated with ErbB signaling pathway and glutamatergic synapse suggesting that LPS suppressed developmental pathways and synapse formation in Purkinje cells (**Figure 3e**). We then performed immunofluorescence for calbindin-1 and measured the Pukinje cell linear density by dividing the number of Purkinje cells by the length of the Purkinje cell folia in 3 cerebellar folia per animal. ^30^ There was a nonsignificant trend towards decreased Purkinje cell linear density in LPS-exposed fetuses consistent with the finding of decrease proportion of Purkinje cells in the single-nuclei analysis (**figure 3e**).

**Figure 3.**
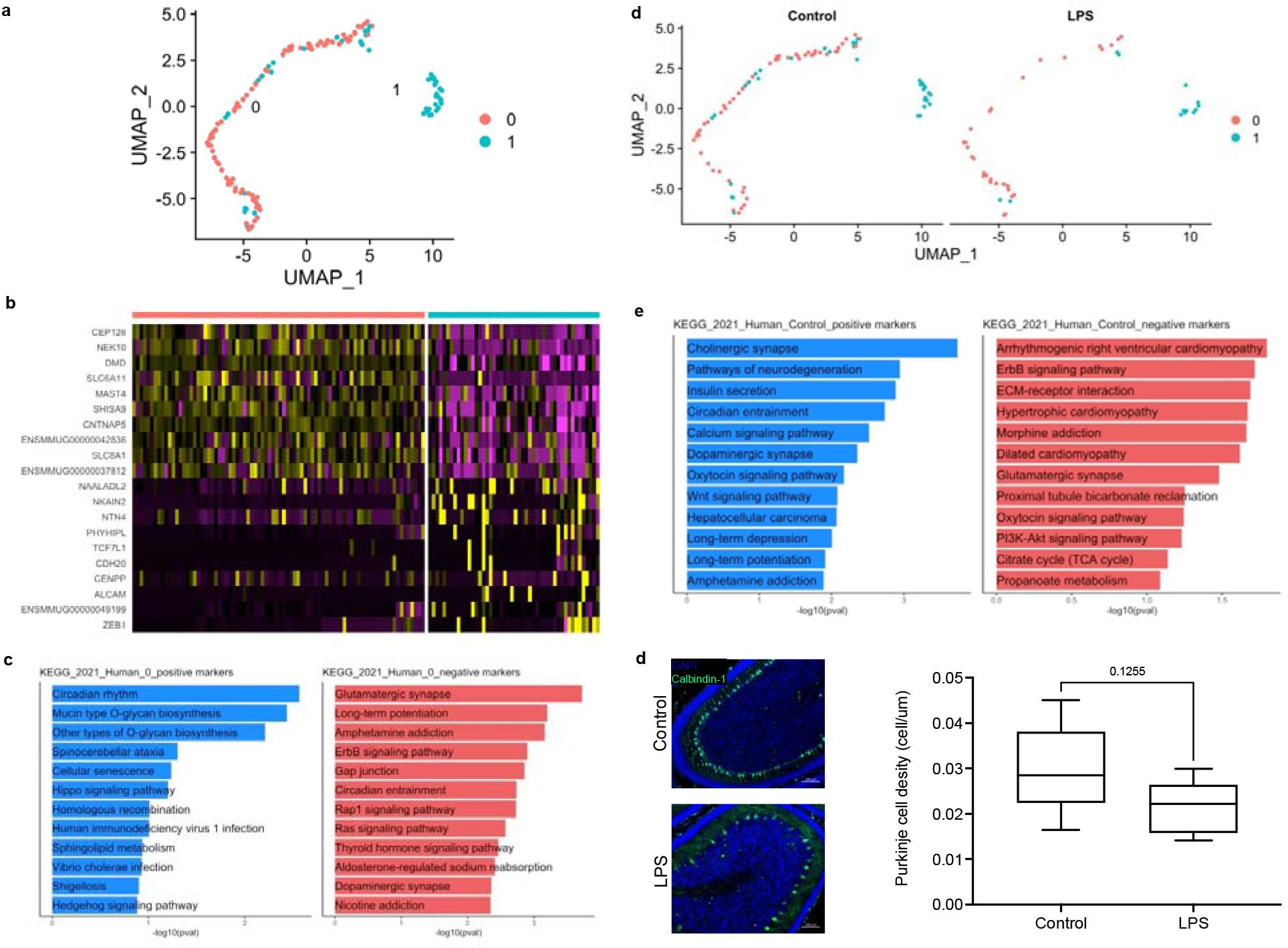
Prenatal inflammation decreases Purkinje cell density in the cerebellum. **a,** UMAP of the cell subset identified as Purkinje cells. Re-clustering of Purkinje cells identified 2 subpopulations. **b,** Hetmap of top differentially expressed gene in each Purkinje cell cluster **c,** Gene set enrichment analysis for Kegg pathways of genes differentially expressed in cluster 1 compared to cluster 0 of Purkinje cells. **d,** UMAP plots of Purkinje cells by condition showing decreased number of Purkinje cells in LPS. **e,** Immunofluorescence (magnification 20X) for Calbindin-1 and Purkinje cell density analysis showing a trend of decreased density of Purkinje cells in LPS-exposed fetuses.

### Chorioamnionitis accelerates oligodendrocyte maturation in the cerebellum

Given the increased proportion of oligodendrocytes in LPS-exposed fetuses relative to control on the single-nuclei data we sought to further determine the effect of chorioamnionitis on oligodendrocyte proliferation and maturation. Sub-setting and re-clustering of oligodendrocytes revealed four distinct clusters (**figure 4a, b**) which were identified based on the top cell markers (**supplemental table 4**) and genes known to be differentially expressed during oligodendrocyte maturation, specifically: Topoisomerase IIα (TOP2A) in proliferating oligodendrocytes, platelet-derived growth factor receptor α (PDGFRA) in oligodendrocyte progenitor cell (OPC), and myelin basic protein (MBP) and myelin-associated glycoprotein (MAG) in premyelinating oligodendrocytes and mature oligodendrocytes (**figure 4a, c**). We then performed differential gene expression analysis in the oligodendrocyte cluster of LPS compared to control fetuses. Overrepresentation analysis of genes differentially expressed between LPS and control revealed that genes suppressed by LPS were associated with oligodendrocyte differentiation and genes induced by chorioamnionitis were associated with myelination (**Figure 4d, supplemental table 7**). We then analyzed cerebellar myelination by Western blotting for MBP, which showed increased MBP in the cerebellum of LPS exposed fetuses (**figure 4e**). To further validate those findings we performed pseudotime analysis on Monocle3^27^ and Slingshot^28^ followed by differential topology between conditions^29^. On both analyses we observed an increased proportion of mature oligodendrocytes in the chorioamnionitis animals (**figure 4f**).

**Figure 4.**
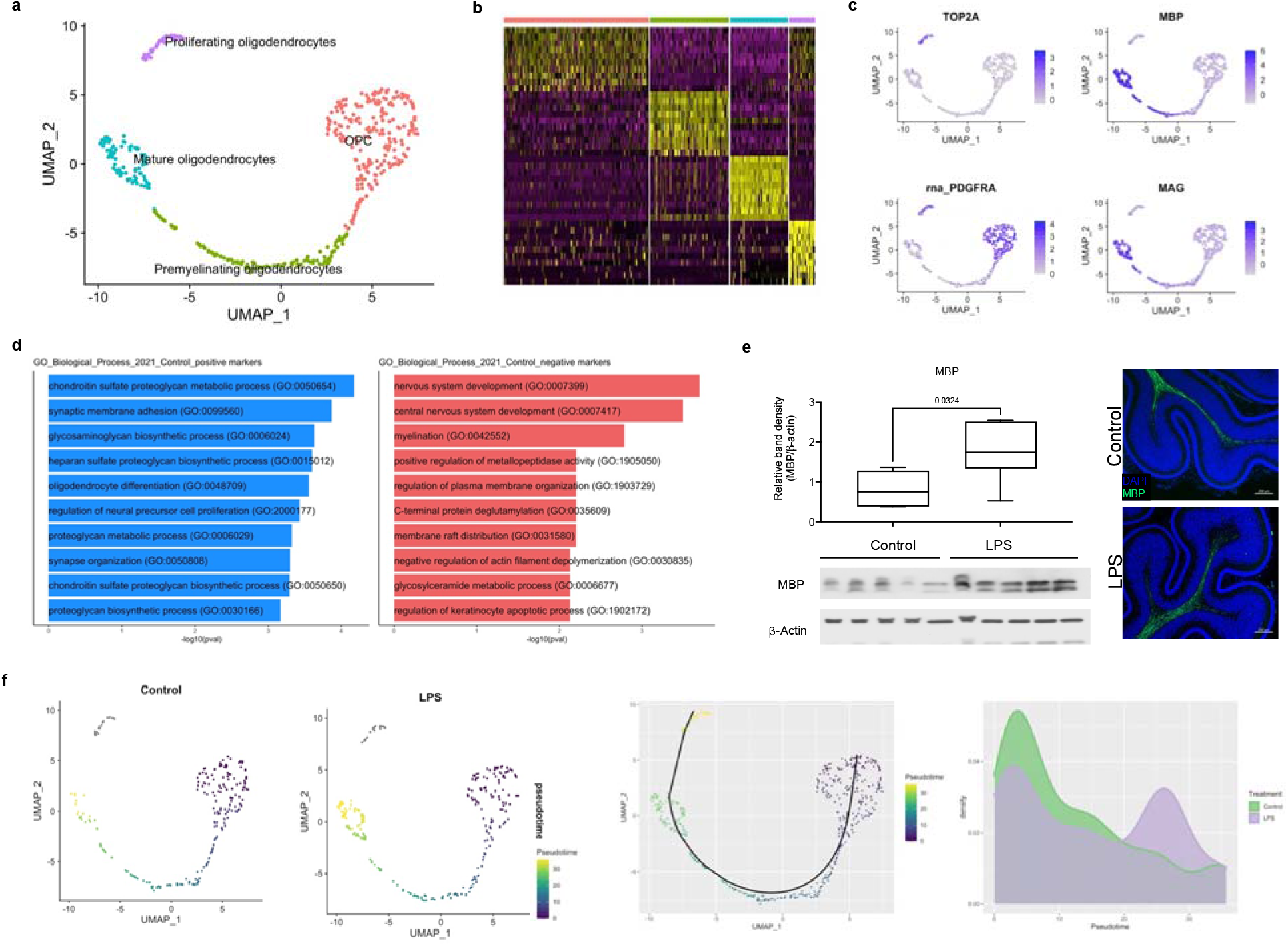
Prenatal inflammation disrupts oligodendrocyte development with increased expression of myelination-associated genes. **a,** UMAP plot of the oligodendrocyte cluster. Re-clustering showed 4 subpopulations of oligodendrocytes that were identified as stages of oligodendrocyte development based on gene expression. **b,** Heatmap of top differentially expressed gene in each oligodendrocyte cell cluster **c,** Expression of markers of oligodendrocyte development in proliferating oligodendrocytes (TOP2A), oligodendrocyte precursor cells (OPC) (PDGFRA), and myelinating / mature oligodendrocytes (MBP and MAG). **d,** Gene set enrichment analysis for Biological processes of genes differentially expressed in the oligodendrocyte cluster in LPS compared to control fetuses with blue as upregulated in control and red as downregulated in control. **e,** Western blot and immunofluorescence for myelin basic protein (MBP) in the fetal cerebellum, p value = 0.03 (n=5 animals/group). **f,** Pseudotime analysis on monocle3 and slingshot using the condiments package to identify differences in proportion of cells along pseudotime shows accelerated oligodendrocyte maturation LPS-exposed fetuses.

### sn-RNAseq reveals microglial and astrocyte diversity in the developing cerebellum

To determine the effect of chorioamnionitis on microglial and astrocyte activation in the cerebellum we individually analyzed the microglia and astrocyte clusters. We found 3 distinct microglia clusters based on expression profile (**supplemental figure 3a, b**). There was no difference in cluster distribution between control and LPS (**supplemental figure 3c**). Top cell markers for microglial clusters (**supplemental table 4**) revealed canonical microglia markers triggering receptor expressed on myeloid cells 2 (TREM2) and CX3CR1 and cluster specific microglial markers, Plexin domain containing 2 (PLXDC2), disable 2 (DAB2), and cell adhesion molecule 2 (CADM2) (**supplemental figure 3d**). Overrepresentation analysis of the top differentially expressed genes in each cluster revealed specific microglial functions. Genes differentially regulated in cluster 0 were associated with mononuclear cell differentiation, microglia migration, and regulation of cell adhesion (**supplemental figure 3e**), cluster 1 was associated with ERK1 and ERK2 cascade, and receptor mediated endocytosis (**supplemental figure 3f**), and cluster 2 was associated with cell adhesion, channel activity, and synapse organization (**supplemental figure 3g**), suggesting cluster specific microglia roles in the developing cerebellum.

Sub-setting and re-clustering of astrocytes revealed 4 distinct cell clusters based on gene expression pattern (**supplemental figure 4a, b**) that showed similar distribution between control and LPS-exposed fetuses (**supplemental figure 4c**). Top cell cluster markers for the four astrocyte clusters identified revealed cluster-specific and cluster enriched gene markers (**supplemental figure 4d supplemental table 4**), which were associated with specific astrocyte biological functions on overrepresentation analysis. Cluster 0 was associated with epidermal growth factor signaling which is crucial for astrocyte transition from quiescent to activated state (**supplemental figure 4e**). Cluster 1 was associated with inositol lipid-mediated signaling, regulation of gene expression, and endothelial proliferation (**supplemental figure 4f**). Cluster 2 was associated with extracellular matrix organization, regulation of synapse potentiation, and axonogenesis (**supplemental figure 4g**). Cluster 3 was associated with synaptic transmission and signaling, dendrite self-avoidance, and glutamate signaling (**supplemental figure 4h**). These findings support the presence of functionally diverse population of astrocytes in the late gestation cerebellum.

### Cell-cell communication in the cerebellum is disturbed by chorioamnionitis

Initial gene differential expression analysis suggested suppression of Purkinje cell-granule cell precursor signaling. To further determine the effect of chorioamnionitis on the intercellular communication networks in the developing cerebellum we analyzed our data on CellChat^31^. CellChat uses established ligand-receptor knowledge for quantitative inference and quantitation of intercellular communication networks. Quantification of the global number of interactions and interaction strength among the cell clusters identified showed that LPS increased the number and strength of inferred interactions among the cell clusters (**Figure 5a, b**). LPS particularly decreased the incoming interaction strength for oligodendrocytes, Purkinje cells, and endothelial mural cells, and increased the incoming interaction strength for proliferating granule cells, Bergman glia, and Golgi cells (**Figure 5c**).

**Figure 5.**
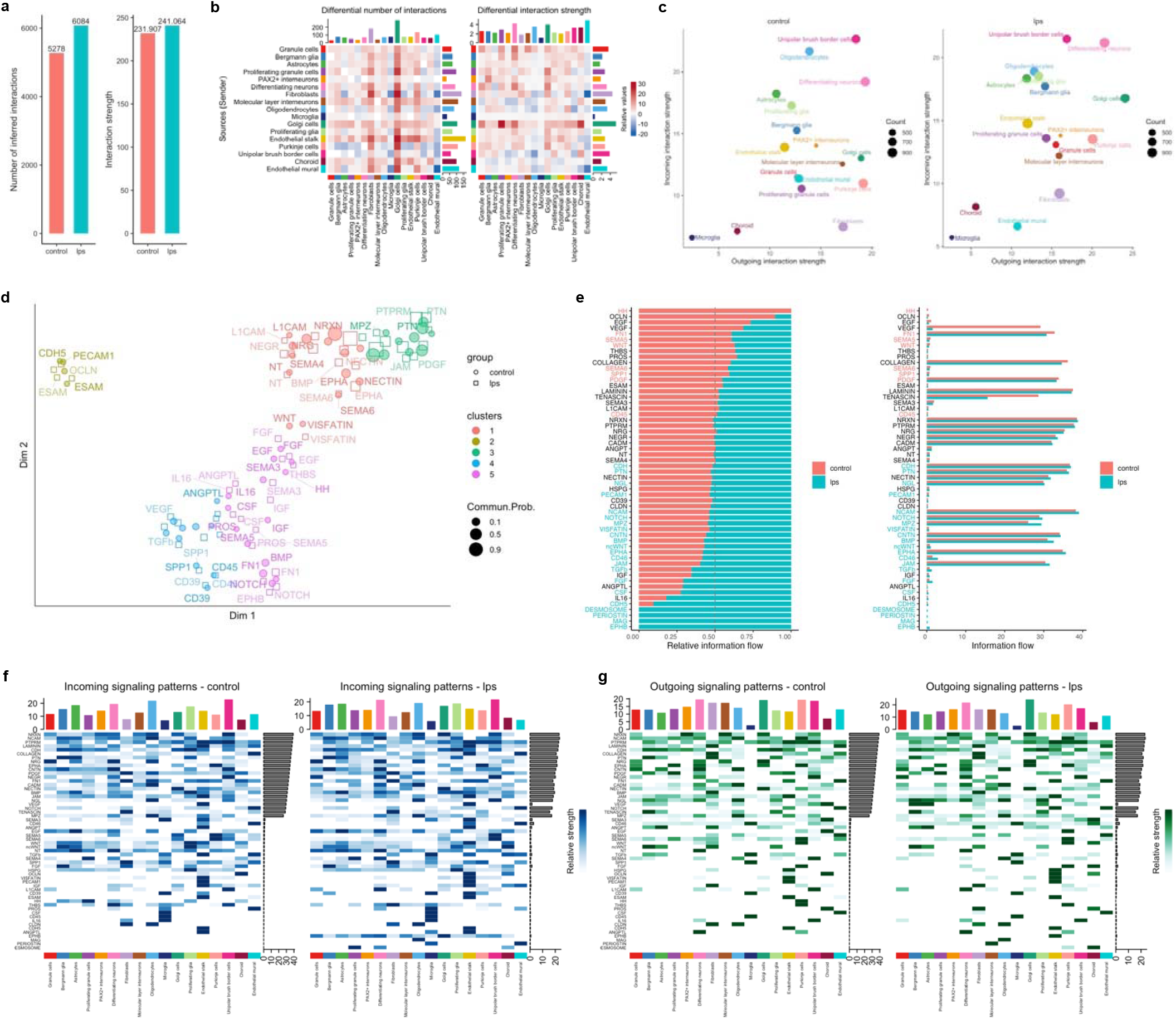
Cell-cell communication is disrupted by antenatal inflammation. **a,** global number of inferred interaction (left) and global strength of interaction (right) in the cerebellum of control and LPS-exposed fetuses. **b,** differential number of interactions and interactions strength between each cell type identified on clustering. Red represents increased, blue represents decreased. The top colored barplot represents the sum of column values displayed in the heatmap (incoming interactions), the right colored barplot represents the sum of row values (outgoing interactions). **c,** scatterplot of outgoing and incoming interaction strength for each cell cluster in two dimensions. **d**, scatterplot and classification of communication networks based on their functional similarity. **e,** barplots of relative (right) and absolute information flow showing conserved and context-specific signaling pathways in control and LPS. **f,** Heatmap of incoming signaling patterns in each cluster in control (right) and LPS (left). **g,** Heatmap of incoming signaling patterns in each cluster in control (right) and LPS (left).

We then investigated how the cell-cell communication architecture changed in chorioamnionitis by projecting the inferred cell communication networks from control and LPS in a shared two-dimensional space based on their functional similarity (**Figure 5d**). We identified 5 major signaling groups based on functional similarity of which some were unique to control such as Hedgehog (HH, cluster 5), and others unique to LPS such as EphrinB (EPHB) and Desmosome in cluster 5, and Periostin and MAG in cluster 4. In addition, other pathways such as WNT, bone morphogenetic protein (BMP), and netrin-F ligand (NGL) were classified into different groups in control and LPS, suggesting that these pathways changed their cell-cell communication architecture in chorioamnionitis (**Figure 5d, supplemental figure 5**).

Next, we determined the conserved and context-specific signaling pathways in control and LPS by comparing the information flow for each signaling pathway (**Figure 5e**). We identified HH as a control-specific pathway, while Desmosome, Periostin, MAG, and EPHB as LPS-specific pathways, albeit with small absolute information flow. Moreover, we identified significant differences in information with both increases in LPS (such as WNT and PDGF) as well as decreases in LPS (such as NOTCH, BMP, and FGF) relative to control. We then analyzed the cell-specific incoming and outcoming signaling patterns for the pathways identified above (**Figure 5f, g**). Purkinje cells were the unique source of HH signaling in controls targeting multiple cell types and HH signaling was not present in LPS. Oligodendrocytes were the unique source and target of MAG in LPS, and MAG signaling was not present in controls.

### Chorioamnionitis impairs SHH signaling from Purkinje cells to proliferating granule cells

Given the finding of suppressed HH signaling from Purkinje cells in LPS-exposed fetuses we further analyzed the differential interactions from Purkinje cells to proliferating granule cells / granule cells between LPS and control. We found several increased inferred interactions in LPS, such as NRG2-ERBB4 which is involved in the formation and maturation of synapses, neural cell adhesion molecule (NCAM) interactions that are involved in the final stages of axonal growth and synaptic stabilization, and FGF-FGFR interactions that inhibit SHH mediated proliferation (**Figure 6a**). Furthermore, we found decreased SHH-PTCH signaling in LPS, which is necessary for maintenance of granule cell proliferation in the external granule layer (**Figure 6a**). Analysis of the expression of the different molecules in the HH pathways revealed decreased expression of SHH in Purkinje cells (**Figure 6b**). Detailed analysis of senders, receivers, and modulators of HH signaling in control animals showed Purkinje cells as the sole sender and proliferating granule cells as the major receiver. We then confirmed localization and expression of SHH at the protein level in the cerebellum (**Figure 6d).** Immunohistochemistry showed SHH localized to Purkinje cell and granule cells with overall decreased signal in LPS. Western blot confirmed decreased SHH in LPS animals relative to control, corroborating that SHH is secreted by Purkinje cells and that SHH production and secretion of SHH is decreased in LPS (**Figure 6d**).

**Figure 6.**
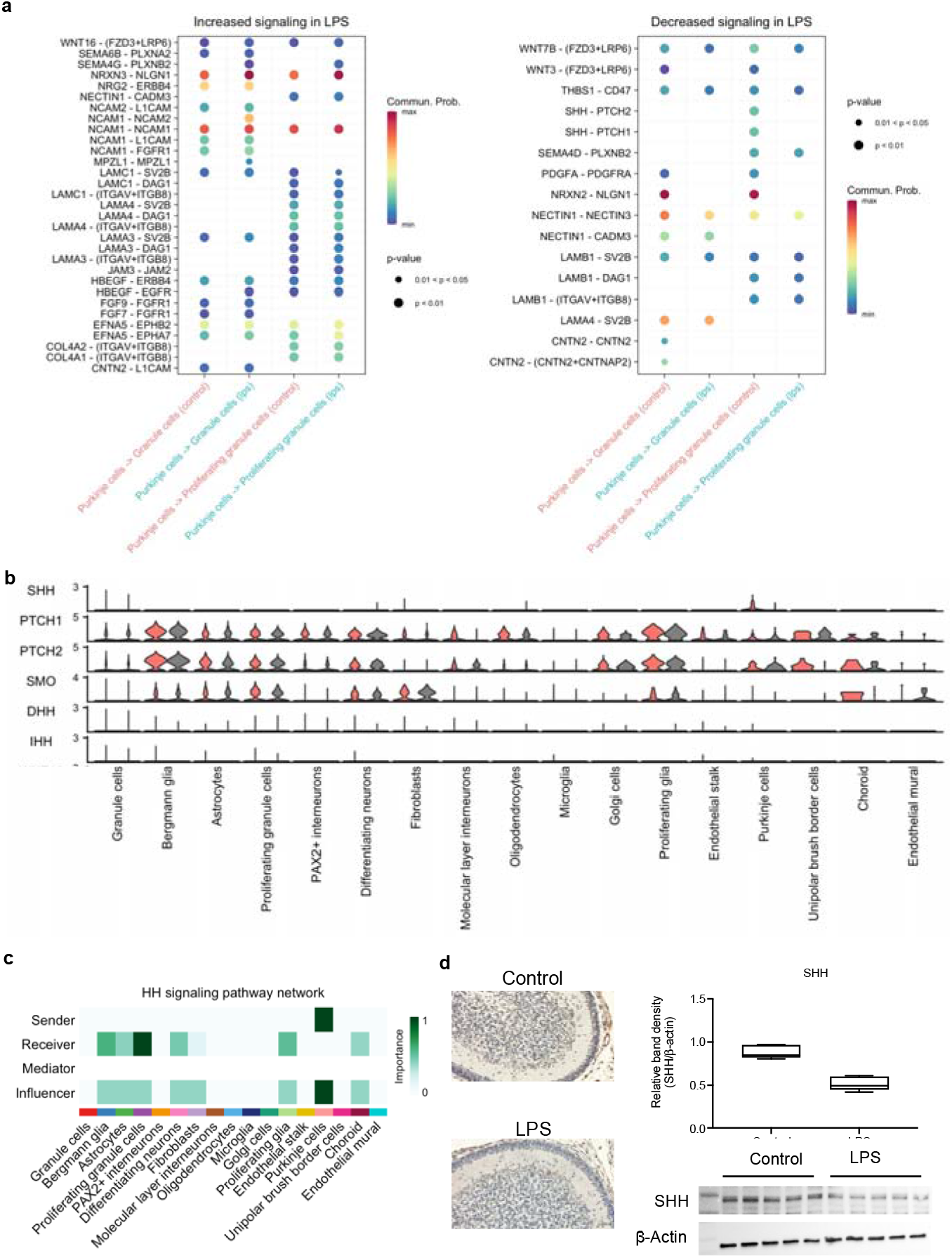
Chorioamnionitis impairs SHH signaling from Purkinje cells into proliferating granule cells. **a,** bubble plots of communication probability of ligand-receptors from Purkinje cells into proliferating granule cells and from Purkinje cells into granule cells upregulated (right) and downregulated (left) in LPS compared to control. **b**, violin plot of signaling genes associated related to SHH signaling. **c**, heatmap of senders, receivers, mediators, and influencers of SHH signaling showing exclusive origin of SHH in Purkinje cells in controls and proliferating granule cells as the dominant receiver. **d**, immunohistochemistry and western blot for SHH showing decreased SHH in the cerebellum of LPS-exposed fetuses compared to controls, p value <0.001 (n=5 animals/group).

### Chorioamnionitis promote maturation signaling in oligodendrocytes

We then delved into the signaling pathways involved in oligodendrocyte differentiation. During transition from oligodendrocyte precursor cell to myelinating oligodendrocyte there is decreased expression of PDGFRA and increased expression of myelin associated genes MBP, MAG, and myelin oligodendrocyte protein (MOG). We found that LPS increased MAG signaling and decreased PDGFC-PDGFRA in oligodendrocytes (**Figure 7a**). Analysis of senders, receivers, and modulators showed that PDGF signaling from proliferating glia, endothelial stalk cells, and UBC was decreased in LPS compared to controls; while MAG signaling was exclusive to LPS-exposed fetuses (**Figure 7b**). At the gene level, chorioamnionitis decreased the expression of PDGFC and PDGFRA and increased the expression of MBP, MAG, and MOG (**Figure 7c**), consistent with our finding of decreased MBP at the protein level (**Figure 4d**).

**Figure 7.**
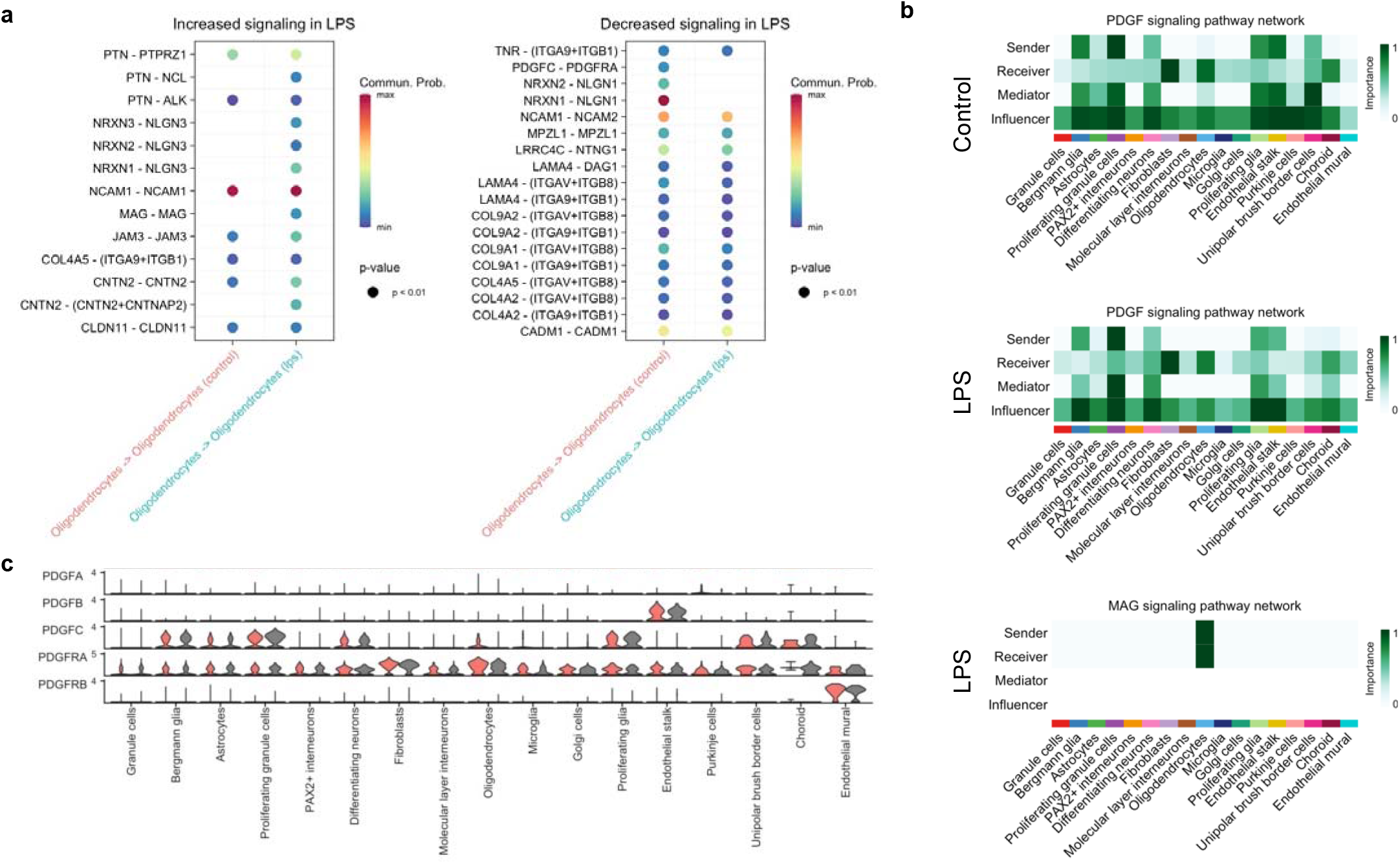
Chorioamnionitis impairs oligodendrocyte maturation signaling. **a,** bubbleplots of communication probability of ligand-receptors from/to oligodendrocytes upregulated (right) and downregulated (left) in LPS compared to control. **b**, heatmap of senders, receivers, mediators, and influencers of PDGF and MAG signaling showing decreased PDGF signaling outgoing and incoming from oligodendrocytes in controls and exclusive MAG signaling in LPS. **c**, violin plot of signaling genes associated related to PDGF-PDGFRA signaling.

## Discussion

In this study we demonstrate, in a clinically relevant model of prematurity and chorioamnionitis using preterm Rhesus macaques, that antenatal inflammation disrupts cerebellar development by impairing hedgehog signaling from Purkinje cells. Impaired hedgehog signaling was associated with accelerated maturation of granule cells and oligodendrocytes on single-nuclei analyses confirmed by translational analyses we show that chorioamnionitis leads to a decrease in the number of Purkinje cells with accelerated maturation of granule cells and oligodendrocytes in the developing cerebellum. We also determine that decreased Purkinje cell-derived SHH signaling to granule cells and pre-oligodendrocytes is driving the disrupted cerebellar development. These findings are consistent with autopsy of individuals with ASD^10^.

The improved survival of extreme preterm infants is obviating the association of preterm birth and ASD. A large cohort study of over 4 million singleton births showed that prevalence of autism in extremely preterm births was as high as 6.1%^32^. A recent metanalysis including 18 studies examining the prevalence of ASD in preterm infants found a similar prevalence rate of 7%^2^. Development of ASD is also linked to prenatal inflammation. The presence of histological chorioamnionitis carries an increased risk of development of ASD^4^. Follow up from a large cohort of preterm infants showed an increased risk for ASD when histological chorioamnionitis was present^33^. The clinical link between antenatal inflammation and neurodevelopmental disorders with ASD-like phenotype is corroborated by animal studies. In mice, maternal immune activation disrupts the integrated stress response in the fetal brain leading to neurobehavioral abnormalities^15^. In pregnant rats injected with intraperitoneal Group B Streptococcus there was histological evidence of chorioamnionitis and the offspring was more likely to show abnormal social interaction, impaired communication, and hyperactivity^34^.

In the preterm rhesus macaque model IA LPS has been shown to induce robust cell neutrophilic recruitment and the chorio-decidual interface, which is associated with systemic fetal inflammatory response ^21,35^. In this model we have observed early neuroinflammation of the cerebellum with increased expression of cytokines and increased IL-6 in the CSF ^20^. At 5 days we found a more subtle but persistent neutrophilic infiltrate and inflammation of the amniotic membranes. The to fetal systemic inflammation induced by IA LPS was associated with decreased number of Purkinje cells by snRNA-seq and a trend of decreased density of Purkinje cells on histology. L of cerebellar Purkinje cells is among the most observed histopathological findings in autism ^10,36,37^. Experimental data from Lurcher mice, which are genetically programmed to have variable degrees of Purkinje cell death after birth, show a correlation between loss of Purkinje cells and deficits on serial reversal-learning task, which measures low order behavioral flexibility^36^.

Purkinje cells are master regulators of the cerebellar development^38^. In rodents, there is a critical window of cerebellar development during the first 2 weeks of life which corresponds to the third trimester cerebellar development in primates. During this period different perinatal insults can lead to Purkinje cell loss and dysfunction. In a rodent model of neonatal brain injury induced by hypoxia there is delayed Purkinje cell arborization and reduction in firing associated with long-term cerebellar learning deficits that can be partially restored by GABA reuptake inhibitors ^39,40^. Exposure to systemic LPS in the second week of life of rodents induces prostaglandin E2 production in the cerebellum and administration of either LPS or prostaglandin E2 at this impairs growth of the Purkinje cell dendritic tree and reduces social play behavior in males, suggesting a developmental-specific role of prostaglandins in cerebellar injury. We have previously shown that IA LPS increases cerebellar COX-2 mRNA in the cerebellum of rhesus macaques at 130 days of gestation^20^. Moreover, genes that regulate Purkinje cell development have been implicated in the pathogenesis of ASD. The Autism susceptibility candidate 2 (AUTS2) was identified as a risk gene for ASD in human genetic studies^41^. Interestingly, AUTS2 is selectively expressed in Purkinje cells and Golgi cells during postnatal development in rodents and conditional deletion of AUTS2 results in an underdeveloped cerebellum with immature Purkinje cells and impaired motor learning and vocal communication^42^.

Purkinje cells regulate cerebellar granule cell maturation by maintaining their proliferative status through SHH signaling^43^. We found that chorioamnionitis led to decreased SHH signaling in Purkinje cells and accelerated maturation of granule cells, with decreased SHH expression in chorioamnionitis confirmed at the protein level. A recent study of preterm baboons without exposure to prenatal inflammation showed that preterm birth was associated with structural and functional changes in Purkinje cells, including decreased synaptic input and abnormal action potentials firing and adaptation^44^. Autopsy of preterm infants showed decreased thickness of the internal and external granule layers with decreased expression of SHH in the Purkinje cell layer compared to term stillborns^19^. Our findings show that prenatal inflammation leads to Purkinje cell loss and disrupted development of granule cells in the developing cerebellum of preterm fetuses that may have important implications to long-term neurodevelopmental outcomes.

In addition to disrupted neuronal maturation, we found that chorioamnionitis also accelerated pre-oligodendrocyte maturation into myelinating oligodendrocytes, evidenced by an increase in myelin-associated genes including MBP and MAG. This finding was consistent with increased expression of MBP on protein analysis. This finding appears to be unique to the cerebellum. A postmortem study from patients with autism showed that in the cerebral white matter myelin-related proteins are decreased, but in the cerebellum they are increased^45^. In addition, a mouse model of placental endocrine deficiency showed a sex-specific effects in cerebellar myelination, with increased myelin in male, which was associated with neurobehavioral abnormalities consistent with autism^46^. While SHH signaling is not a classic mechanism of oligodendrocyte maturation, in vitro studies with cerebellar organotypic cultures have shown that SHH produced by Purkinje cells stimulates the proliferation of oligodendrocyte precursor cells and the decreased SHH production during postnatal development is associated with maturation of oligodendrocytes in the cerebellum^46^. This data suggests that the loss of Purkinje cells with decreased SHH signaling in LPS-exposed fetuses may be a common mechanism to accelerated granule cell and oligodendrocyte maturation.

Overall, our findings in a preterm nonhuman primate model of chorioamnionitis support the role of prenatal inflammation in disrupted cerebellar development through reduction of Purkinje cells associated with accelerated maturation of granule cells and oligodendrocytes. It is well established the link between preterm birth and development of ASD, but the specific mechanisms are unclear. Our results suggest that prenatal inflammation, a common occurrence, and cause of preterm birth, contributes to the disrupted cerebellar development leading to changes similar to the histopathological findings of ASD in the cerebellum. The mechanisms of chorioamnionitis leading to disrupted cerebellar development and ASD may be common to other inflammatory exposure during pregnancy including congenital infections and viral illnesses such as influenza.

## Methods

### Animal experiments

Animal protocols were reviewed and approved by the IACUC and the University of California Davis. Time-mated pregnant rhesus macaques at 127 days of gestation (80% of term gestation) were treated with intra-amniotic LPS 1mg (*E. coli* O55:B5, Sigma-Aldrich) diluted in 1 mL of sterile saline (LPS). Controls received no intervention. Animals were delivered *en cul* by cesarean section at 5 days after intra-amniotic LPS (Figure 1A). No spontaneous labor or fetal losses were observed in the intervention or control groups. Fetuses were euthanized with pentobarbital. After euthanasia the cerebellum was dissected and the hemispheres were separated along the midline of the vermis. The left hemisphere was frozen for molecular analysis and the right hemisphere was fixed in 10% formalin and embedded in paraffin for histological analyses. Data of animals included in the study as show in **supplemental table 1**.

### Chorion-amnion-decida dissection and flow cytometry of chorio-decidua cells

At delivery, extra-placental membranes were collected and dissected away from the placenta as previously described ^35^. The cells of the chorio-decidua were scraped and the amnion and remaining chorion tissue were separated with a forceps. The chorio-decidua cells were washed and digested with Dispase II (Life Technologies, Gran Island, NY) plus collagenase A (Roche, Indianapolis, IN) followed by DNase I (Roche) treatment. Cells suspensions were filtered, red blood cells lysed, and the suspension was prepared for flow cytometry. Cell viability was >90% by trypan blue exclusion test. Multiparameter flow cytometry and gating strategy of the difference leukocyte subpopulations was done as previously described ^35^. Monoclonal antibodies used are listed in **supplemental table 8**. Cells were treated with 20 μg/mL of human immunoglobulin G (IgG) to block Fc receptors, stained for surface markers for 30 min at 4°C in PBS, washed, and fixed in fixative stabilizing buffer (BD Bioscience). All antibodies were titrated for optimal detection of positive populations and similar mean fluorescence intensity. At least 500,000 events were recorded for each sample. Doublets were excluded based on forward scatter properties, and dead cells were excluded using LIVE/DEAD Fixable Aqua dead cell stain (Life Technologies). Unstained and negative biological population were used to determine positive staining for each marker. Data were analyzed using FlowJo version 9.5.2 software (TreeStar Inc., Ashland, OR).

### Cytokine concentration measurement

We measured cytokine concentration in maternal and fetal plasma by Luminex technology using multiplex kits for nonhuman primate (Millipore). Concentrations were calculated from standard curves for recombinant proteins.

### Immunofluorescence and immunohistochemistry

Paraffin-embedded tissue sections underwent heat-assisted antigen retrieval with citrate buffer (pH 6.0). For immunohistochemistry endogenous peroxidase activity was reduced with CH3OH/H2O2 treatment. Nonspecific binding sites were blocked with 4% bovine serum diluted in PBS followed by incubation with primary antibodies overnight at 4 °C (**supplemental table 9**). The following day, tissue sections were incubated with the appropriate species-specific biotinylated or Alexa Fluor-conjugated secondary antibody diluted in 1:200 in 4% bovine serum for 2 hours at room temperature. For immunohistochemistry with DAB, tissue sections incubated with biotinylated secondary antibody were washed and antigen/antibody complexes were visualized using Vectastain ABS peroxidase kit (Vector Laboratories, Burlingame, CA) followed by counterstaining with Harris hematoxylin. For immunofluorescence, sections incubated with Alexa Fluor-conjugated secondary antibody were washed and mounted with VectaShield Hardset Mounting Media (Vector Laboratories).

### Statistical analyses

GraphPad Prism (GraphPad Software, La Jolla, California, USA) was used to graph and analyze data. Statistical differences between groups were analyzed using Mann-Whitney U-tests for cytokine concentration measurements and cellular composition from flow cytometry results. Results were considered significantly different for p values ≤0.05. However, due to the limited number of samples per group, we also report trends (p values between 0.05 and 0.1).

### Single-nucleus RNA isolation, sequencing, and mapping

Nuclei isolation and snRNA sequencing (sn-RNAseq) were performed by Singulomics Corporation (Singulomics.com). Flash frozen cerebellar hemispheres from 2 rhesus macaque fetuses per group were homogenized and lysed with Triton X-100 in RNase-free water for nuclei isolation. Isolated nuclei were purified, centrifuged, and resuspended in PBS with RNase inhibitor, and diluted to 700 nuclei/μl for standardized 10x capture and library preparation protocol using 10x Genomics Chromium Next GEM 3’ Single Cell Reagent kits v3.1 (10x Genomics, Pleasanton, CA). The libraries were sequenced with an Illumina NovaSeq 6000 (Illumina, San Diego, CA). Raw sequencing files were processed with CellRanger 5.0 (10x Genomics) and mapped to the rhesus macaque genome (Mmul_8.0.1) with the option include introns and generated count matrices.

### SnRNA-seq processing, clustering, and differential expression

Downstream analyses were performed on the Seurat package version 3.1.0 for R^47^. Each sample was processed separately and then integrated for cell type identification and comparison analyses. After importing the CellRanger data using the Read10x function and creating a Seurat object for each sample, samples were normalized, scaled, and the top 2000 variable features were identified followed by nuclei clustering. After clustering, ambient RNA was removed with SoupX^48^. Cleaned samples were then filtered removing nuclei with <500 genes or >5% mitochondrial RNA. Number of nuclei and genes per samples is shown in **supplemental table 2**. After filtering, samples were integrated using the FindIntegrationAnchors function with the parameter dims = 1:30 followed by IntegrateData function with dims = 1:30. The integrated data was scaled, principal component analysis (PCA) was performed with npcs = 30, and nuclei were clustered with resolution = 0.5. Conserved cell types for each cluster were identified using the FindAllMarkers function with default parameters. Cell types were identified based on known cell markers and previously published datasets of cerebellum single-cell sequencing ^22–24^. Differences in the proportion of cells in clusters between control and IA LPS was performed using the permutation test on the scProportionTest package for R^25^. Global and cluster subset differential expression analyses between Control and IA LPS were performed using the FindMarkers function with default parameters. Overrepresentation analysis of differentially expressed genes was performed for Gene Ontology terms and KEGG Pathways on ToppCluster^26^ and using the DEenrichRPlot function on Seurat^49,50^.

### Pseudotime analyses

To determine lineage trajectories and determine differences across conditions we performed pseudotime analyses with Monocle3 ^27,51,52^ and Slingshot^28^ followed by *condiments*^29^. For analysis on Monocle3^27^, *celldataset* objects were created from the individual Seurat objects and combined using the combine_cds function and preprocessed. We then performed dimensionality reduction using UMAP ^53^ followed by cell clustering. Specific cell types of interest were subset, clustered and a principal graph was fit within each partition using the learn_graph function. Cells were then ordered in pseudotime using the plot_cells function. For analysis with Slingshot^28^ a Seurat object for each condition was created and converted to a *single cell experiment* object followed by conversion to slingshot object using the slingshot function with reducedDim = “PCA”. Objects were hen filtered and normalized following the standard protocol. We then performed the topologyTest on condiments^29^ to determine differences in lineage trajectories between conditions.

### Cell communication analyses

Cell-cell interactions were inferred with CellChat^31^ based on known ligand-receptor pairs in difference cell types. To identify perturbed cell-cell communication networks in chorioamnionitis we loaded the condition specific Seurat objects into cell chat using the createCellChat function, followed by the preprocessing functions indentifyOverEpressedGenes, identifyOverExpressedinteractions, and projectData with default settings. For comparison between Control and IA LPS, we applied the computeCommunProb, computeCommunProbPathway, and aggregateNet functions using standard parameters and fixed randomization seeds. To determine signal senders and receivers we used the netAnalysis_signalingRole function on the netP data slot.

### Availability of data and materials

The gene expression data have been deposited in NCBI’s Gene Expression Omnibus (GEO) and are accessible through GEO Series accession no. GSE205001.

## Author contributions

JN and AFS designed the sequencing and molecular experiments, performed the snRNA-seq analysis interpreted the data and wrote the manuscript with input from the other authors. JN, XT, AT, VNP, PSK, KW, AD, MTN performed the molecular and histological experiments and analysis of the data. PP performed the flow cytometry and ELISA experiments. MB, SW, KY and RB, contributed the data analysis and interpretation. LAM, SFK, CAC, AHJ, AFS conceived and supervised the animals experiments.

---

**Supplemental figure 1.**
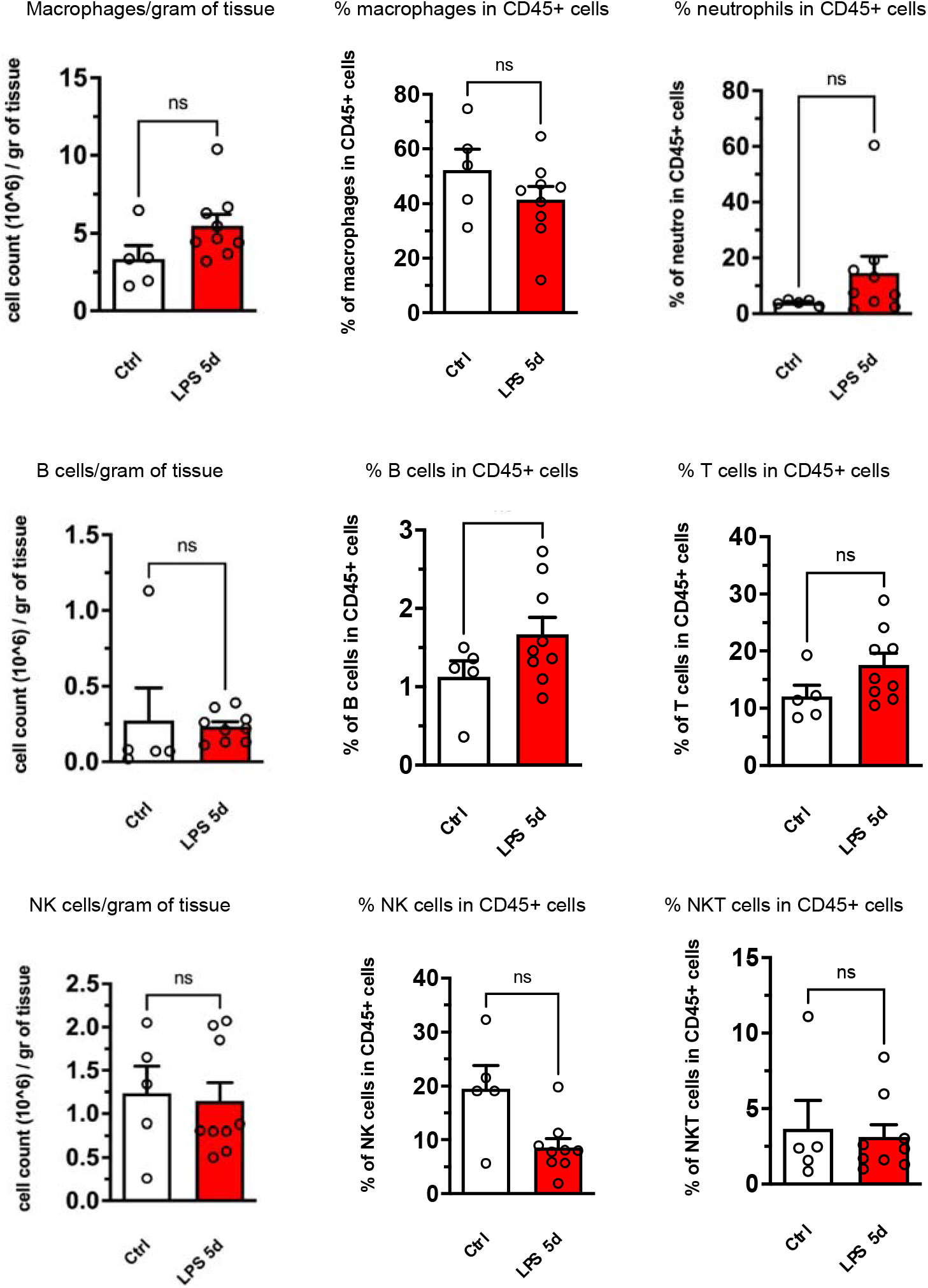
Flow cytometry of the chorio-decidua cells.

**Supplemental figure 2.**
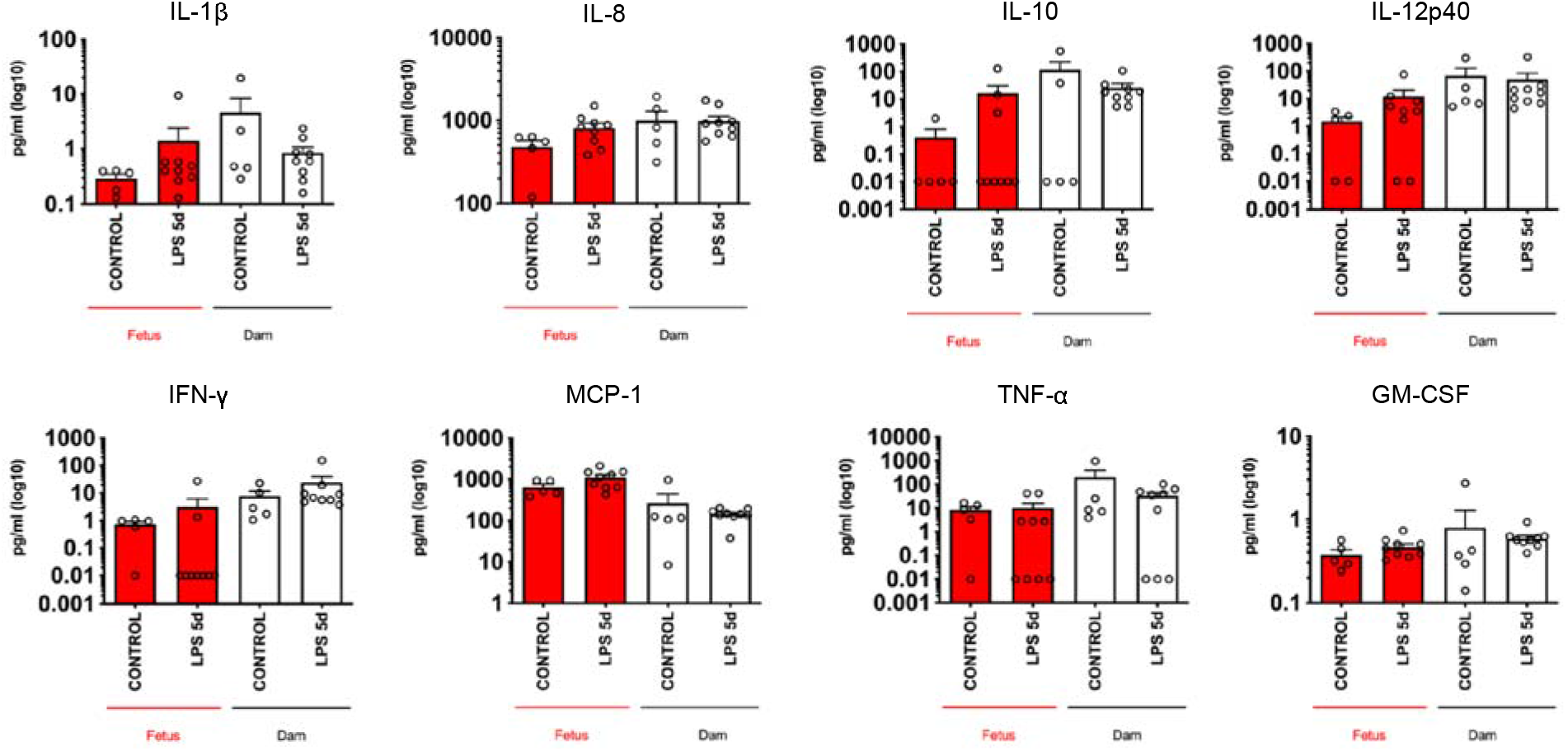
Multiplex ELISA for cytokines in the maternal and fetal plasma.

**Supplemental figure 3.**
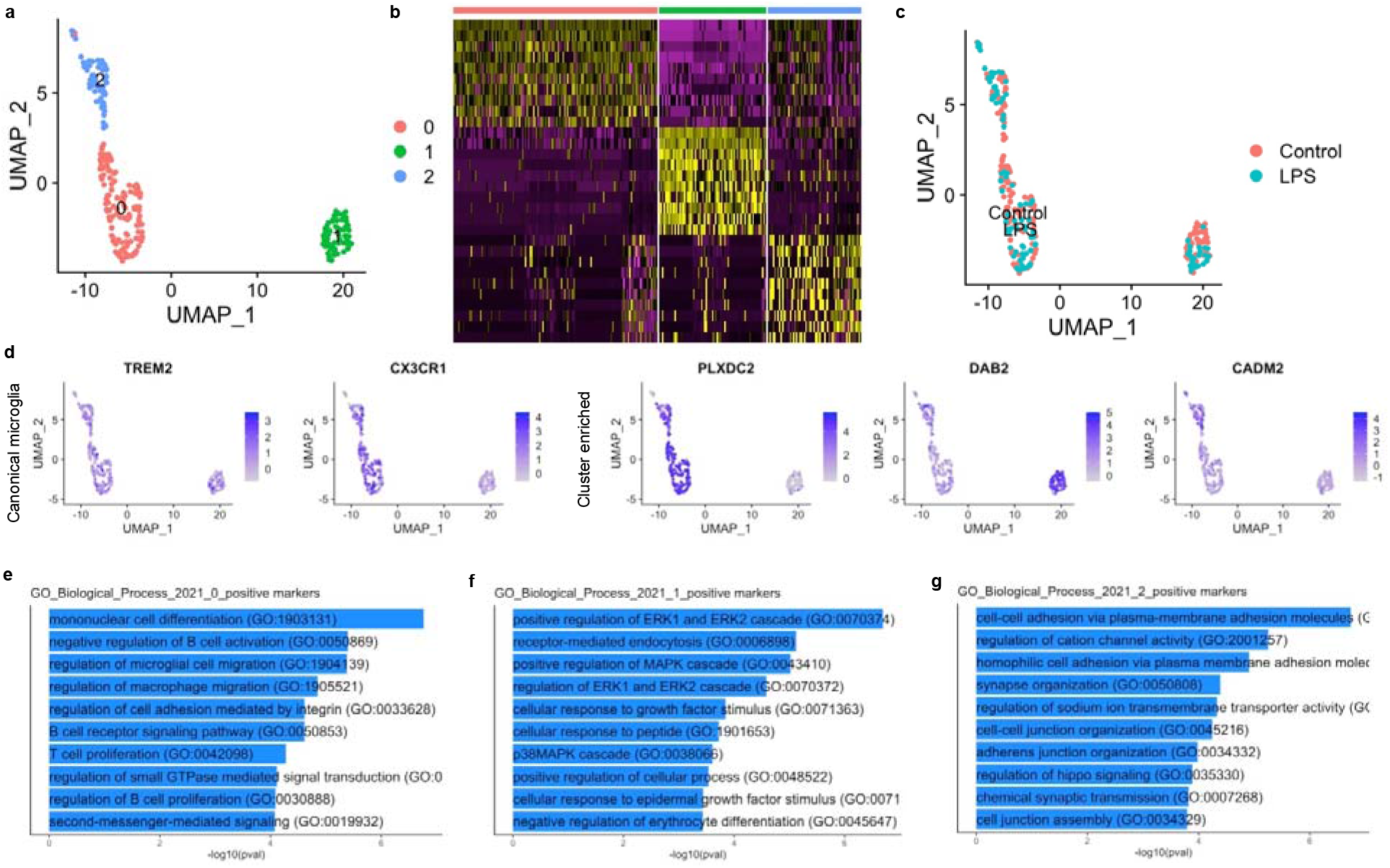
Microglial diversity in the developing cerebellum. **a,** UMAP plot of the microglia cluster showing 3 subpopulations. **b,** Heatmap of top differentially expressed genes in each microglia cluster **c,** UMAP plot of microglia cluster by condition. **d,** Expression of canonical and cluster-specific microglial markers. **e**, Gene set enrichment analysis for Biological processes of genes differentially regulated in cluster 0. **f**, Gene set enrichment analysis for Biological processes of genes differentially regulated in cluster 1. **g**, Gene set enrichment analysis for Biological processes of genes differentially regulated in cluster 2.

**Supplemental figure 4.**
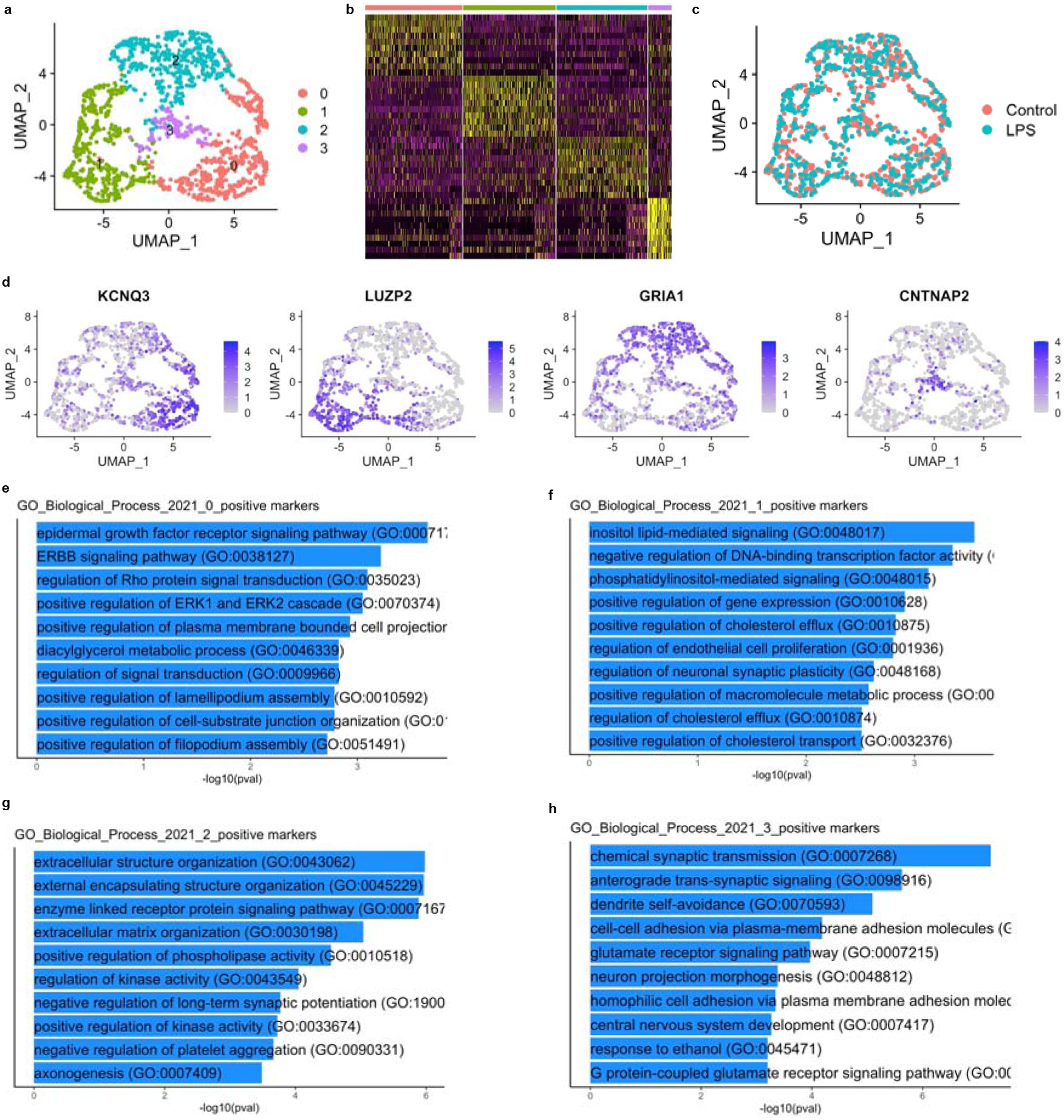
Astrocyte diversity in the developing cerebellum. **a,** UMAP plot of the astrocyte cluster showing 3 subpopulations. **b,** Heatmap of top differentially expressed genes in each astrocyte cluster **c,** UMAP plot of astrocyte clusters by condition. **d,** Expression of cluster-specific markers. **e**, Gene set enrichment analysis for Biological processes of genes differentially regulated in cluster 0. **f**, Gene set enrichment analysis for Biological processes of genes differentially regulated in cluster 1. **g**, Gene set enrichment analysis for Biological processes of genes differentially regulated in cluster 2. **h**, Gene set enrichment analysis for Biological processes of genes differentially regulated in cluster 3.

**Supplemental figure 5.**
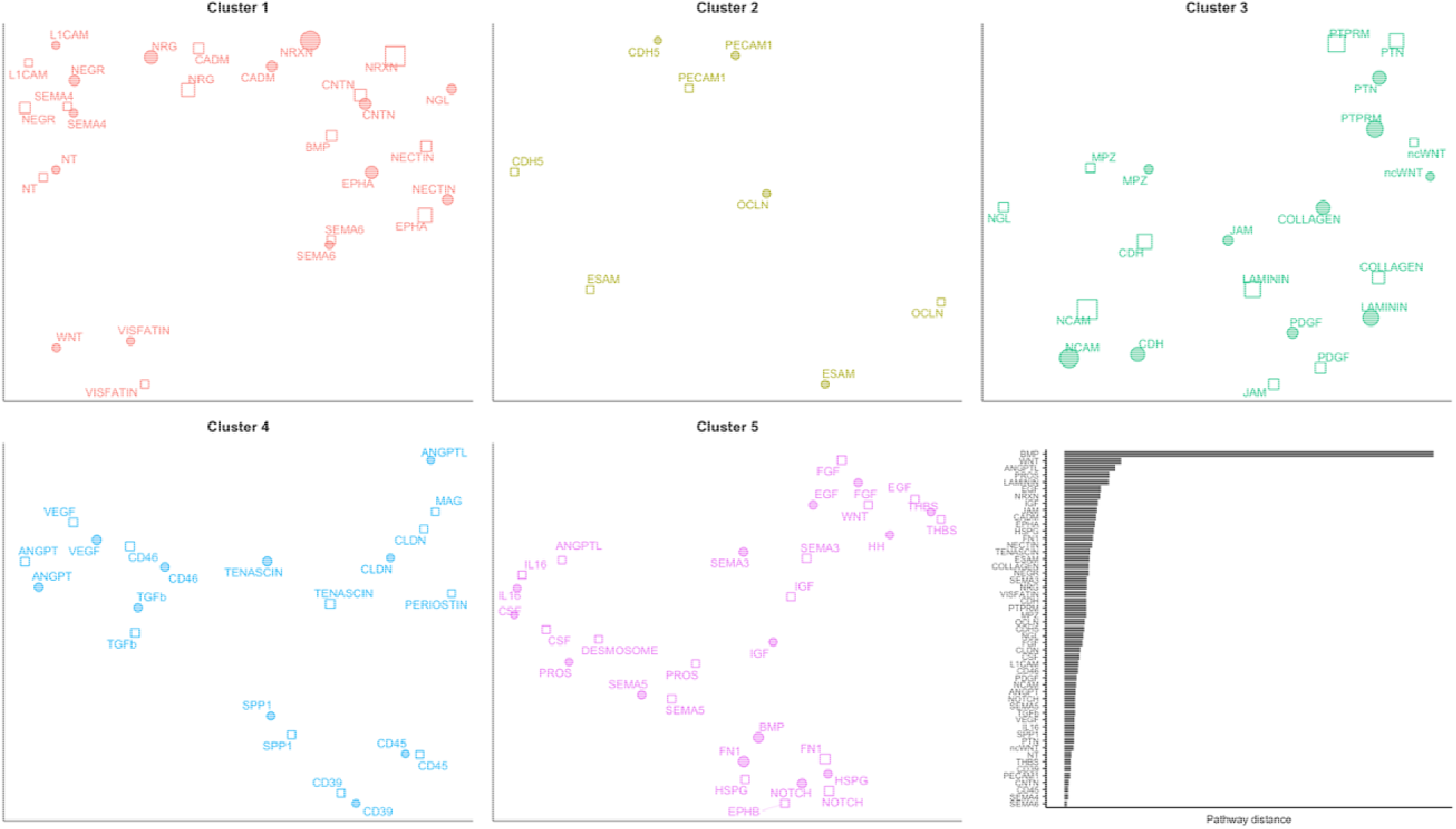
Classification of cellular communication networks based on function similarity.

**Supplemental table 1.** Characteristics of animals included in the study.

**Supplemental table 2.** Characteristics of samples included in snRNA-seq.

**Supplemental table 3.** Top 10 differentially expressed genes in each of the 24 clusters identified, used for cell type identification.

**Supplemental table 4.** Top 10 differentially expressed genes for specific cell clusters: granule cells, Purkinje cells, oligodendrocytes, microglia, and astrocytes.

**Supplemental table 5.** Differentially expressed genes in LPS compared to controls in the granule cell cluster.

**Supplemental table 6.** Differentially expressed genes in LPS compared to controls in the Purkinje cell cluster.

**Supplemental table 7.** Differentially expressed genes in LPS compared to controls in the oligodendrocyte cluster.

**Supplemental table 8.** Antibodies used in flow cytometry experiments.

**Supplemental table 9.** Antibodies used in flow immunohistochemistry and western blots.

