## supplemental table 1 for "Single-nucleus RNA-seq reveals disrupted cell maturation by chorioamnionitis in the preterm cerebellum of nonhuman primates"

Supplementary table 1. Characteristics of animals included in the study

| | **Control (n= 8)** | **IA LPS (n=9)** |
|---|---|---|
| Gestational age at delivery (days median, range) | 133 (130-135) | 133 (131-134) |
| Birth weight (g mean, SD) | 330 ± 35 | 321 ± 37 |
| Gender (F/M) | 2/6 | 4/5 |
| Amniotic fluid cell count ($10^4$/mL median, range) | 0 (0-40) | 30 (22-60) * |
| Maternal plasma WBC ($10^9$/L average, SD) | 8.2 ± 2.3 | 5.6 ± 1.7 * |
| Maternal plasma neutrophils (% average, SD) | 78 ± 6 | 51 ± 16 * |
| Maternal plasma lymphocytes (% average, SD) | 18 ± 5 | 43 ± 15 * |
| Maternal plasma platelets ($10^9$/L average, SD) | 356 ± 71 | 349 ± 88 |
| Fetal plasma WBC ($10^9$/L average, SD) | 2.4 ± 0.6 | 3.3 ± 0.8 * |
| Fetal plasma neutrophils (% average, SD) | 9.7 ± 6 | 23 ± 5 * |
| Fetal plasma lymphocytes (% average, SD) | 85 ± 8 | 73 ± 6 * |
| Fetal plasma platelets ($10^9$/L average, SD) | 369 ± 72 | 314 ± 91 * |

WBC: white blood cell count; SD: standard deviation; $^*p<0.05$.
