## supplemental table 2 for "Single-nucleus RNA-seq reveals disrupted cell maturation by chorioamnionitis in the preterm cerebellum of nonhuman primates"

Supplementary table 2. Characteristics of samples included in snRNA-seq.

| | **Control 1** | **Control 2** | **LPS 1** | **LPS 2** |
|---|---|---|---|---|
| Gestational age at delivery | 130 | 130 | 131 | 133 |
| Birth weight (g) | 290 | 290 | 282 | 348 |
| Gender | F | M | F | M |
| Number of nuclei | 9,021 | 8,593 | 7,002 | 6,095 |
| Mean reads per nuclei | 25,192 | 30,437 | 24,008 | 45,124 |
| Median genes per nuclei | 2,118 | 1,203 | 1,990 | 1,589 |
