## supplemental table 8 for "Single-nucleus RNA-seq reveals disrupted cell maturation by chorioamnionitis in the preterm cerebellum of nonhuman primates"

Supplementary table 8. Monoclonal antibodies used in the flow cytometry of chorio-decidua cells.

| **Antibody** | **Clone** | **Conjugation** | **Manufacturer** |
|---|---|---|---|
| CD45 | D058-1283 | PE-CF594 | BDBioscience |
| HLA-DR | L243 | Brilliant Violet 570<br>PercPCy5.5 | Biolegend |
| CD3 | SP34-2 | APC-Cy7 | BDBioscience |
| CD14 | TUK4 | Pacific Blue | Thermofisher |
| CD56 | NCAM16.2 | PE-Cy7 | BDBioscience |
| CD88 | P12/1 | Alexa Fluor 647 | AbD Serotec |
| CD19 | HIB19 | Alexa Fluor 700 | Biolegend |
| CD20 | 2H7 | Alexa Fluor 700 | Biolegend |
| CD16 | 3G8 | Alexa Fluor 700 | BDBioscience |
| CD63 | H5C6 | Pacific Blue | Biolegend |
| Live/Dead | | Aqua | BDBioscience |
