## supplemental table 9 for "Single-nucleus RNA-seq reveals disrupted cell maturation by chorioamnionitis in the preterm cerebellum of nonhuman primates"

Supplementary table 9. Antibodies used in the Western blots, immunohistochemistry, and immunofluorescence experiments.

| **Antibody** | **Host** | **Application** | **Dilution** | **Catalog** | **Manufacturer** |
|---|---|---|---|---|---|
| Beta-actin | Mouse | WB | 1:10000 | A5441 | Sigma-Aldrich |
| Calbindin | Rabbit MAb | IF | 1:200 | Ab108404 | Abcam |
| MBP | Mouse MAb | IF | 1:3000 | ab62631 | Abcam |
| MBP | Mouse MAb | WB | 1:10000 | ab62631 | Abcam |
| SHH | Mouse MAb IgG1 | IHC | 1:50 | ab135240 | Abcam |
| SHH | Mouse MAb IgG1 | WB | 1:1000 | ab135240 | Abcam |
| Anti-Mouse IgG | Goat | WB | 1:10000 | 31430 | ThermoFisher |
| Anti-rabbit AF488 | Goat | IF | 1:200 | A32731 | ThermoFisher |
| Anti-rabbit IgG | Goat | WB | 1:10000 | 31460 | ThermoFisher |
| Anti-rabbit IgG | Goat | IHC | 1:200 | BA1000 | Vector Labs |

IHC: immunohistochemistry; IF: immunofluorescence, MAb: monoclonal antibody, MBP: myelin basic protein, SHH: sonic hedgehog, WB: Western blotting
